# Compensatory evolution drives multidrug-resistant tuberculosis in Central Asia

**DOI:** 10.1101/334599

**Authors:** Matthias Merker, Maxime Barbier, Helen Cox, Jean-Philippe Rasigade, Silke Feuerriegel, Thomas A. Kohl, Roland Diel, Sonia Borrell, Sebastien Gagneux, Vladyslav Nikolayevskyy, Sönke Andres, Ulrich Nübel, Philip Supply, Thierry Wirth, Stefan Niemann

## Abstract

Bacterial factors favoring the unprecedented multidrug-resistant tuberculosis (MDR-TB) epidemic in the former Soviet Union remain unclear.

We utilized whole genome sequencing and Bayesian statistics to analyze the evolutionary history, temporal emergence of resistance and transmission networks of MDR-MTBC strains from Karakalpakstan, Uzbekistan (2001-2006).

One MTBC-clone (termed Central Asian outbreak, CAO) with resistance mediating mutations to eight anti-TB drugs existed prior the worldwide introduction of standardized WHO-endorsed directly observed treatment, short-course (DOTS). DOTS implementation in Karakalpakstan in 1998 likely selected for these CAO-strains, comprising 75% of sampled MDR-TB strains in 2005/2006. CAO-strains were also identified in a previously published cohort from Samara, Russia (2008-2010). Similarly, transmission success and resistance development was linked to mutations compensating fitness deficits associated with rifampicin resistance.

The genetic make-up of these outbreak clades threatens the success of both empirical and standardized guideline driven MDR-TB therapies, including the newly WHO-endorsed short MDR-TB regimen in Uzbekistan.

## Introduction

Multidrug-resistant tuberculosis (MDR-TB), caused by *Mycobacterium tuberculosis* complex (MTBC) strains that are resistant to the first-line drugs isoniazid and rifampicin, represent a threat to global TB control. Barely 20% of the estimated annual 480,000 new MDR-TB patients have access to adequate second-line treatment regimens. The majority of undiagnosed or ineffectively treated MDR-TB patients continue to transmit their infection and suffer high mortality (1).

Based on early observations that the acquisition of drug resistance could lead to reduced bacterial fitness (2) it was hypothesized that drug-resistant MTBC-strains had a reduced capacity to transmit, and would not widely disseminate in the general population (3–7). This optimistic scenario has been invalidated by the now abundant evidence for transmission of MDR and extensively drug-resistant MTBC-strains (XDR-TB; MDR-TB additionally resistant to at least one fluoroquinolone and one aminoglycoside) in healthcare and community settings (3, 8–11). In former Soviet Union countries, which experience the highest MDR-TB rates worldwide, the expansion of drug-resistant MTBC-clones is thought be promoted by interrupted drug supplies, inadequate implementation of regimens, lack of infection control and erratic treatment in prison settings (12, 13). Continued transmission is thought to be aided by the co-selection of mutations in the bacterial population that compensate for a fitness cost (e.g. growth deficit) associated particularly with the acquisition of rifampicin resistance mediating mutations (3, 7–11). The compensatory mechanism for rifampicin resistant MTBC strains is proposed to be associated with structural changes in the RNA-polymerase subunits *RpoA, RpoB*, and *RpoC* that increase transcriptional activity and as a consequence enhance the growth rate (11). However, the impact of these bacterial genetic factors on the epidemiological success of MDR-MTBC strains and implications for current and upcoming MDR-TB treatment strategies remain unexplored.

We utilized whole genome sequencing (WGS) to retrace the longitudinal transmission and evolution of MTBC-strains towards MDR/pre-XDR/XDR geno- and phenotypes in Karakalpakstan, Uzbekistan. In this high MDR-TB incidence setting, the proportion of MDR-TB among new TB-patients increased from 13% in 2001 to 23% in 2014 despite the local introduction of the World Health Organization recommended DOTS strategy in 1998 and an initially limited MDR-TB treatment program in 2003 (14, 15). We expanded our analyses by including a WGS dataset of MDR-MTBC strains from Samara, Russia (2008-2010) (13) to investigate clonal relatedness, resistance and compensatory evolution in both settings.

## Results

### Study population and MTBC phenotypic resistance (Karakalpakstan, Uzbekistan)

Despite differences in sampling for cohort 1 and cohort 2 (see methods), patients showed similar age, sex distributions, and proportion of residence in Nukus, the main city in Karakalpakstan (Uzbekistan) (Appendix Table S1). While the majority of strains from both cohorts were phenotypically resistant to additional first-line TB drugs (i.e. beyond rifampicin and isoniazid), combined resistance to all 5 first-line drugs was significantly greater in cohort 2 (47% in cohort 2 compared to 14% in cohort 1, *P* < 0.0001). The same was true for resistance to the second-line injectable drug capreomycin (23% in cohort 2 compared to 2% in cohort 1, *P* = 0.0001) (Appendix Table S1). This finding was surprising as the isolates from cohort 2 patients - who were treated with individualized second-line regimens predominately comprising ofloxacin as the fluoroquinolone and capreomycin as the second-line injectable - were all obtained before the initiation of their treatment. In addition, there was no formal MDR-TB treatment program in Karakalpakstan prior to 2003. These elements imply that the higher rate of resistance to capreomycin was attributable to infection by already resistant strains (i.e. to primary resistance).

### MTBC population structure and transmission rates

Utilizing WGS, we determined 6,979 single nucleotide polymorphisms (SNPs) plus 537 variants located in 28 genes and upstream regions associated with drug resistance and bacterial fitness (additional data). The corresponding phylogeny revealed a dominant subgroup comprising 173/277 (62.5%) closely related strains within the Beijing-genotype (alias MTBC lineage 2) (Fig. 1). This group, termed Central Asian Outbreak (CAO), showed a highly restricted genetic diversity (median pairwise distance of 21 SNPs, IQR 13-25) and was differentiated from a set of more diverse strains by 38 specific SNPs (Appendix Figure S1, additional data). The proportion of CAO-strains was similar between 2001-02 and 2003-04 (49% and 52% respectively), but increased to 76% in 2005-06 (*P* < 0.01). Over the same time periods, the proportions of other strain types remained stable or decreased (Appendix Figure S2).

**Fig. 1:**
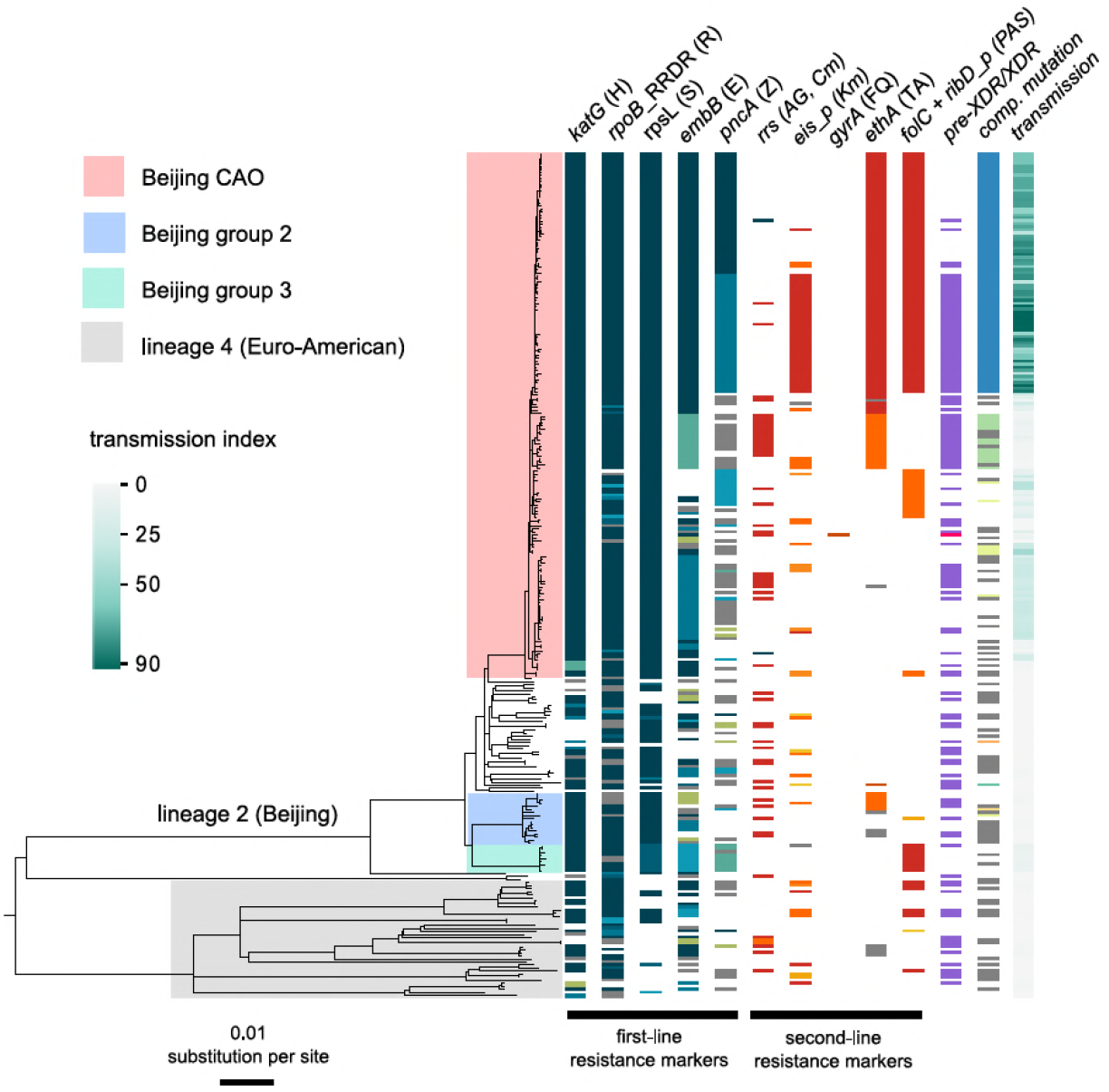
Drug resistance and transmission success among MDR-MTBC strains from Karakalpakstan, Uzbekistan. Maximum likelihood phylogeny (GTR substitution model, 1,000 resamples) of 277 MDR-MTBC strains from **Karakalpakstan**, Uzbekistan sampled from 2001 to 2006. Columns show drug resistance associated mutations to first- and second-line drugs (different mutations represented by different colors), genetic classification of pre-XDR (purple) and XDR (pink) strains, and putative compensatory mutations in the RNA polymerase genes *rpoA, rpoB* and *rpoC*. Transmission index represents number of isolates within a maximum range of 10 SNPs at whole genome level. MTBC-Beijing strains (lineage 2) are differentiated into three sub-groups (i.e. Central Asian Outbreak (CAO), group 2 and group 3). Strains belonging to lineage 4 (Euro-American) are colored in grey: H=isoniazid, R=rifampicin, S=streptomycin, E=ethambutol, Z=pyrazinamide, FQ=fluoroquinolone, AG=aminoglycosides, Km=kanamycin Cm=capreomycin, TA=thioamide, PAS=para-aminosalicylic acid.

We then sized transmission networks (measured by transmission indexes, see methods) supposed to reflect human-to-human transmission over the last ∼10 years based on a maximum of 10 differentiating SNPs between two strains. Transmission rates varied, even among closely related outbreak strains (Fig. 1). Beijing-CAO-strains formed particularly large transmission networks (>50 strains/patients; Fig. 1); 96.0% (166/173) of all Beijing-CAO strains were associated with recent transmission (i.e. transmission index ≥1), versus 48.4% (31/64) of non-CAO Beijing strains (*P*<0.0001) and 57.5% (23/40) of non-Beijing strains (*P*<0.0001) (additional data). In addition the large CAO transmission network exhibited higher levels of drug resistance relative to non-Beijing strains, as reflected by the larger number of drugs for which phenotypic (*P*=0.0079) and genotypic drug resistance (*P*=0.0048) was detected Appendix Figure S3).

### Evolutionary history of CAO strains in Karakalpakstan

In order to gain more detailed insights into the emergence of resistance mutations in the evolutionary history of the CAO clade, we sought to employ a Bayesian phylogenetic analysis for a temporal calibration of the CAO phylogeny and an estimation of the mutation rate. Using an extended collection of more diverse CAO strains (n=220) from different settings (see methods) we initially compensated for the restricted sampling time frame of the Karakalpakstan dataset (2001-2006). A linear regression analysis showed correlation between sampling year and root-to-tip distance and a moderate temporal signal (*P*=0.00039, R^2^= 5.2%, Appendix Figure S4), allowing for further estimation of CAO mutation rates and evaluation of molecular clock models using Bayesian statistics. Based on the marginal L estimates collected by path sampling, we found a strict molecular clock with tip dates to be most appropriate (Appendix Table S2). Mutation rate estimates (under a relaxed clock model) ranged on average from 0.88 to 0.96 × 10^−7^ substitutions per site per year (s/s/y), depending on the demographic model, in favor for the Bayesian skyline model with mutation rate of 0.94 × 10^−7^ (s/s/y) (95% HPD 0.72-1.15 × 10^−7^ (s/s/y)) (Appendix Table S2).

We then employed the Bayesian skyline model with a strict molecular clock set to 0.94 × 10^−7^ (s/s/y) specifically for the CAO clade from Karakalpakstan (n=173). We determined that the most recent common ancestor (MRCA) of the CAO-clade emerged around 1976 (95% highest posterior density (HPD) 1969- 1982). The MRCA already exhibited a streptomycin resistance mutation (*rpsL* K43R) (Fig. 2), and acquired isoniazid resistance (*katG* S315T) in 1977 (95% HPD 1973-1983). The CAO-population size then rose contemporaneously with multiple events of rifampicin, ethambutol, ethionamide, and para-aminosalicylic acid resistance acquisition in different branches (Fig. 2). As an illustration, the most frequent CAO-clone (upper clade in Fig. 2) acquired ethambutol and ethionamide resistance mutations (*embB* M306V, *ethA* T314I) around 1984 (95% HPD 1982-1989), and an MDR-genotype (*rpoB* S450L) around 1986 (95% HPD 1985-1992). The effective population size reached a plateau before fixation of mutations in the *ribD* promoter region (leading to para-aminosalicylic acid resistance) and *rpoC* N698S, putatively enhancing its fitness around 1990 (95% HPD 1989-1994) (Fig. 2). Independent fixation of pyrazinamide (*pncA* Q10P and I133T) and kanamycin (*eis* −12 g/a) resistance-associated mutations was detected in 1992 and 1991 (both with 95% HPD rounded to 1991-1996) (Fig. 2).

**Fig. 2:**
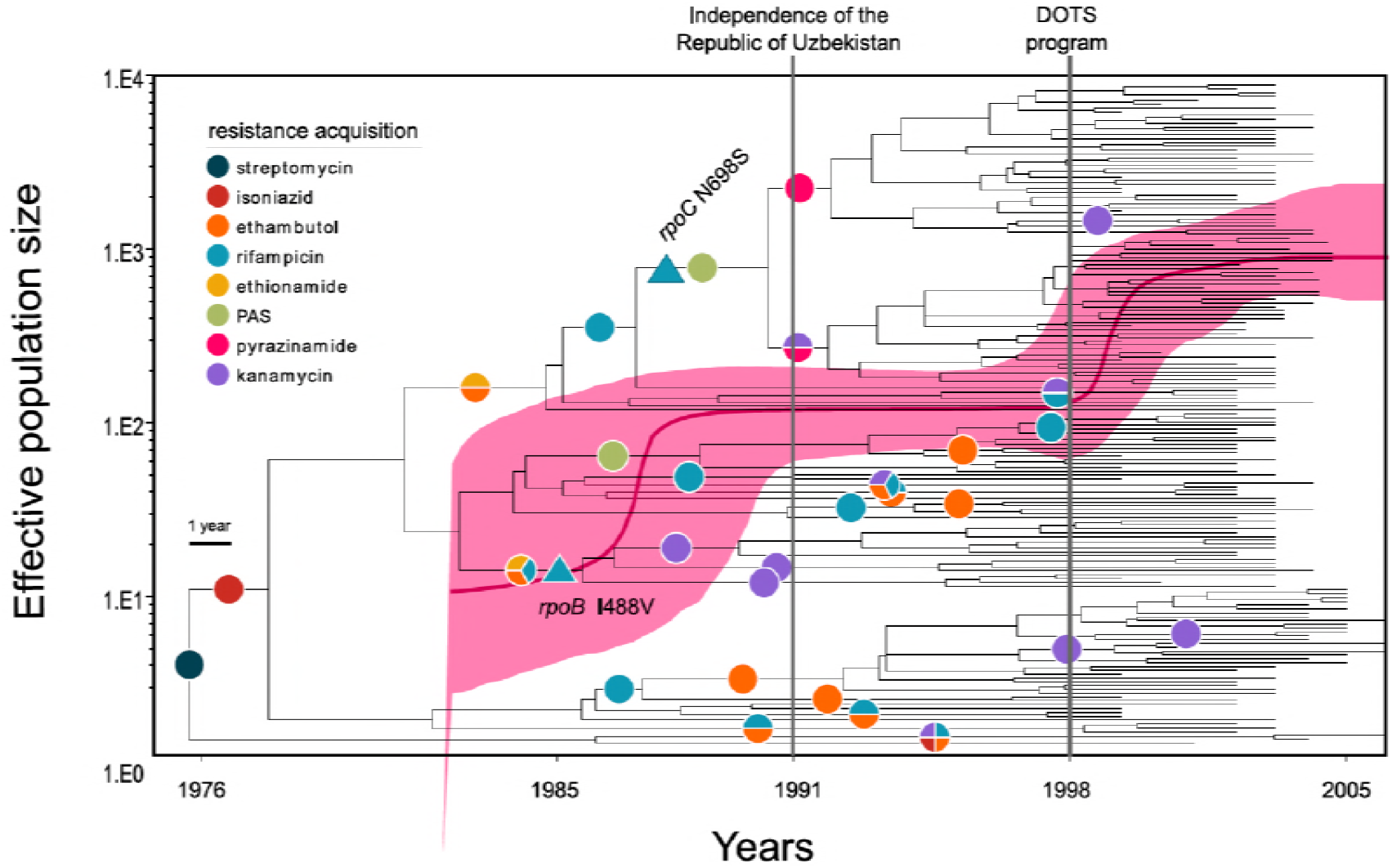
Evolutionary history of MTBC Central Asian outbreak (CAO) strains. Genealogical tree of CAO strains in **Karakalpakstan**, Uzbekistan and effective population size over time based on a (piecewise-constant) Bayesian skyline approach using the GTR substitution model and a strict molecular clock prior of 0.94 × 10^−7^ substitutions per nucleotide per year. Pink shaded area represents changes in the effective population size giving the 95% highest posterior density (HPD) interval with the pink line representing the mean value. Vertical lines indicate time points of the implementation of the first standardized TB treatment program (DOTS) in Karakalpakstan and of the declaration of Uzbekistan as independent republic. Symbols on branches show steps of fixation of resistance conferring mutations.

Interestingly, the implementation of the systematic DOTS-program in Karakalpakstan in 1998 coincided with a second effective population size increase (Fig. 2). At that time, distinct CAO-subgroups already exhibited pre-XDR (in this context MDR plus kanamycin resistance) resistance profiles, mediating resistance to as many as eight different anti-TB drugs. Of note, only a single strain was identified as harboring a *gyrA* mutation (A90V), associated to fluoroquinolone resistance (additional data). At the end of the study period in 2006 we observed a pre-XDR rate among CAO strains of 52.0% (90/173), compared to 35.9% (23/64) among other Beijing strains (*P*=0.03) and compared to 42.5% (17/40) among non-Beijing strains (*P*=0.30) (additional data).

### Impact of compensatory variants on transmission networks

Overall, 62.1% (172/277) of all MDR-MTBC strains carried putative compensatory mutations (11, 13) in *rpoA* (n=7), *rpoC* (n=126) and *rpoB* (n=43) (additional data). These mutations were almost completely mutually exclusive, as only 4/172 strains harbored variants in more than one RNA polymerase-encoding gene. While mutations in *rpoA* and *rpoB* were equally distributed between Beijing-CAO strains and other non-outbreak Beijing strains, CAO-strains had more *rpoC* variants (56% vs 28%, *P*=0.003) (Appendix Table S3). The mean number of resistance mutations was higher among strains carrying compensatory mutations (Fig. 3A), 4.77 vs 3.35 mutations (two-sample t-test *P*=1.2×10^−10^). Notably, strains with compensatory mutations also showed larger transmission indexes than strains presenting no compensatory mutation, 37.16 vs 9.22 (Welch two-sample t-test *P*<2.2×10^−16^) (Fig. 3B). CAO-strains with compensatory mutation also had more resistance-conferring mutations than CAO-strains lacking such mutation (ANOVA, Tukey multiple comparisons of means *P* adj=0.0000012). There was no difference observed for the means of resistance-conferring mutations amon non-CAO strains; compensatory mutation present vs. absent (*P* adj= 0.1978623) (Fig. 3C).

**Fig. 3:**
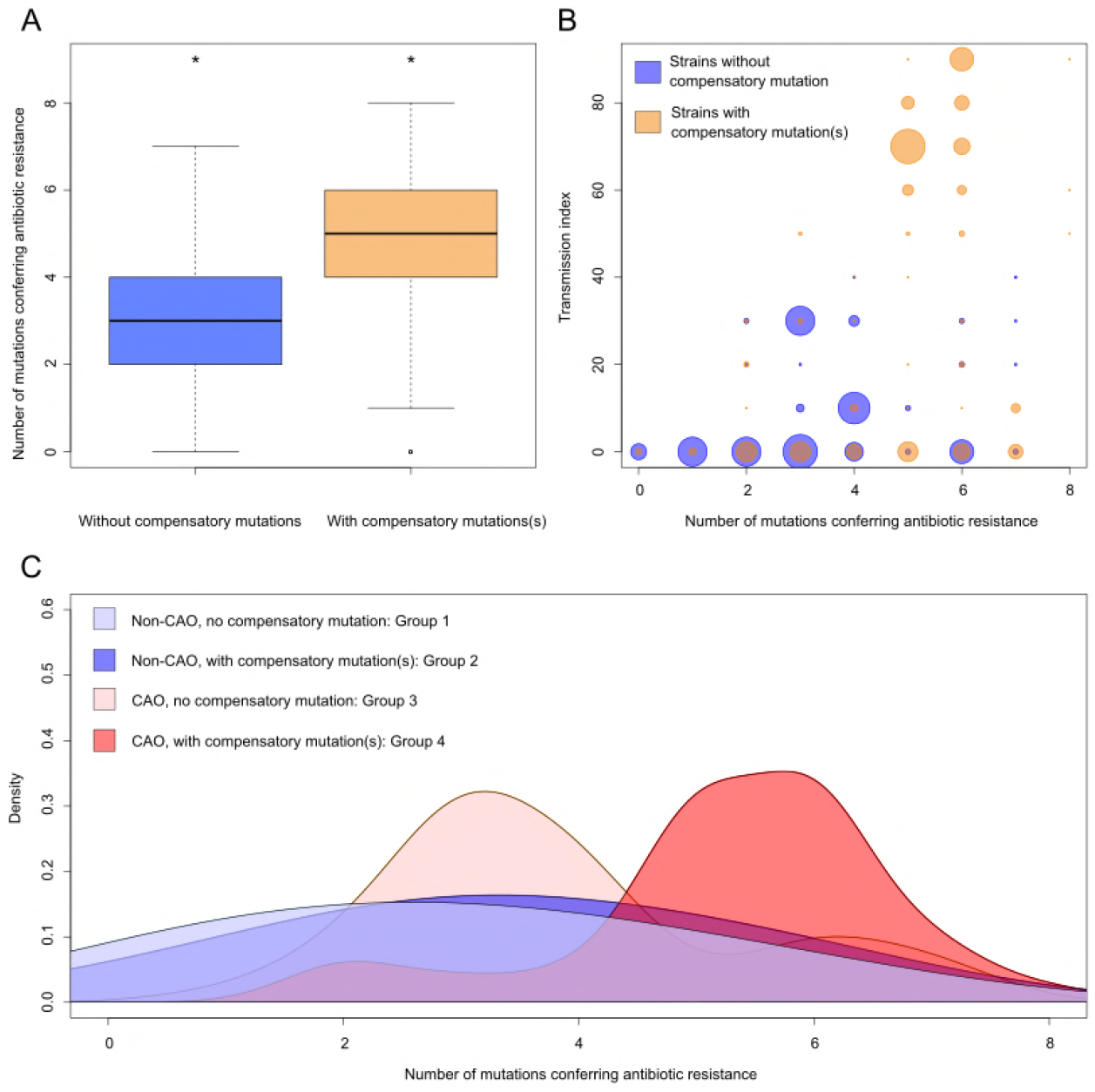
Compensatory mutations and drug resistance levels. Comparisons between strains carrying compensatory mutations (in orange) and strains with no-compensatory mutations (in blue), from the **Karakalpakstan** dataset. A) Boxplot showing number of resistance mutations for the two categories (without or with compensatory mutations). The two categories were significantly different (two-sample t-test *P*=1.2×10^−10^). B) Bubble plots showing the transmission index (number of strains differing by less than 10 SNPs) as a function of antibiotic resistance related mutations. Bubble sizes are proportional to the numbers of strains. C) Density plot of the number of resistance-conferring mutations for 4 groups of strains sourced from the Karakalpakstan data. Proportions are adjusted by using Gaussian smoothing kernels. The 4 groups are composed of non-CAO strains with no compensatory mutations; non-CAO strains carrying compensatory mutations; CAO strains with no compensatory mutations and CAO strains carrying compensatory mutations. These groups are respectively colored in light blue, dark blue, light orange and light red.

Regression-based analyses of transmission success scores in the Beijing-CAO clade confirmed that the presence of compensatory mutations was strongly associated with cluster sizes independent of the accumulation of resistance mutations (Fig. 4). This pattern was mostly observed for clusters initiated in the late 1980s and the 1990s.

**Fig. 4:**
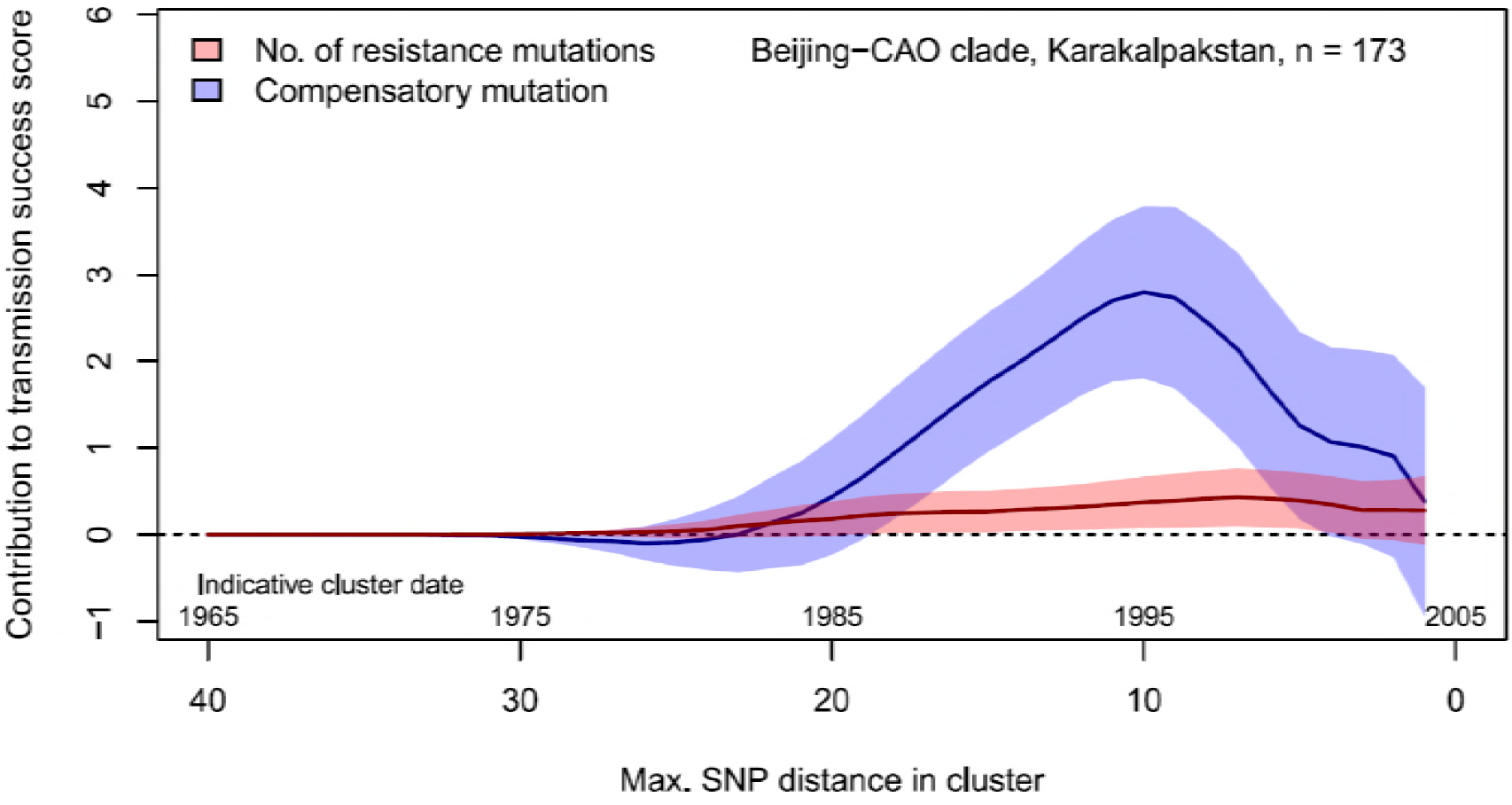
Contributions of resistance-conferring and compensatory mutations to the transmission success of *M. tuberculosis* of the Beijing-CAO clade, **Karakalpakstan**, Uzbekistan. Shown are the coefficients and 95% confidence bands of multiple linear regression of the transmission success score, defined as the size of clusters diverging by at most *N* SNPs and divided by *N* or, equivalently, the size of clusters that evolved over *N* years divided by *N*. The presence of compensatory mutations was independently associated with transmission success, with a maximum association strength found for SNP distances ranging from 10 to 20 SNPs, corresponding to transmission clusters beginning around 1995.

### Combined analysis of MDR-TB cohorts from Karakalpakstan and Samara (2001-2010)

To place our analyses in a broader phylogenetic and geographic context, we combined our Karakalpakstan genome set with previously published genomes of 428 MDR-MTBC isolates from Samara (13), a Russian region located ˜1,700 km from Nukus, Karakalpakstan. This analysis showed that Beijing-CAO strains accounted for the third largest strain clade in Samara (13). Conversely, the second largest clade in Samara, termed Beijing clade B according to Casali et al (13, 21), or European/Russian W148 (22), was represented in Karakalpakstan by a minor clade (Fig. 5). Considering a third Beijing clade (termed clade A) restricted to Samara (13), three major Beijing outbreak clades accounted for 69.6% (491/705) of the MDR-TB cases in both regions.

**Fig. 5:**
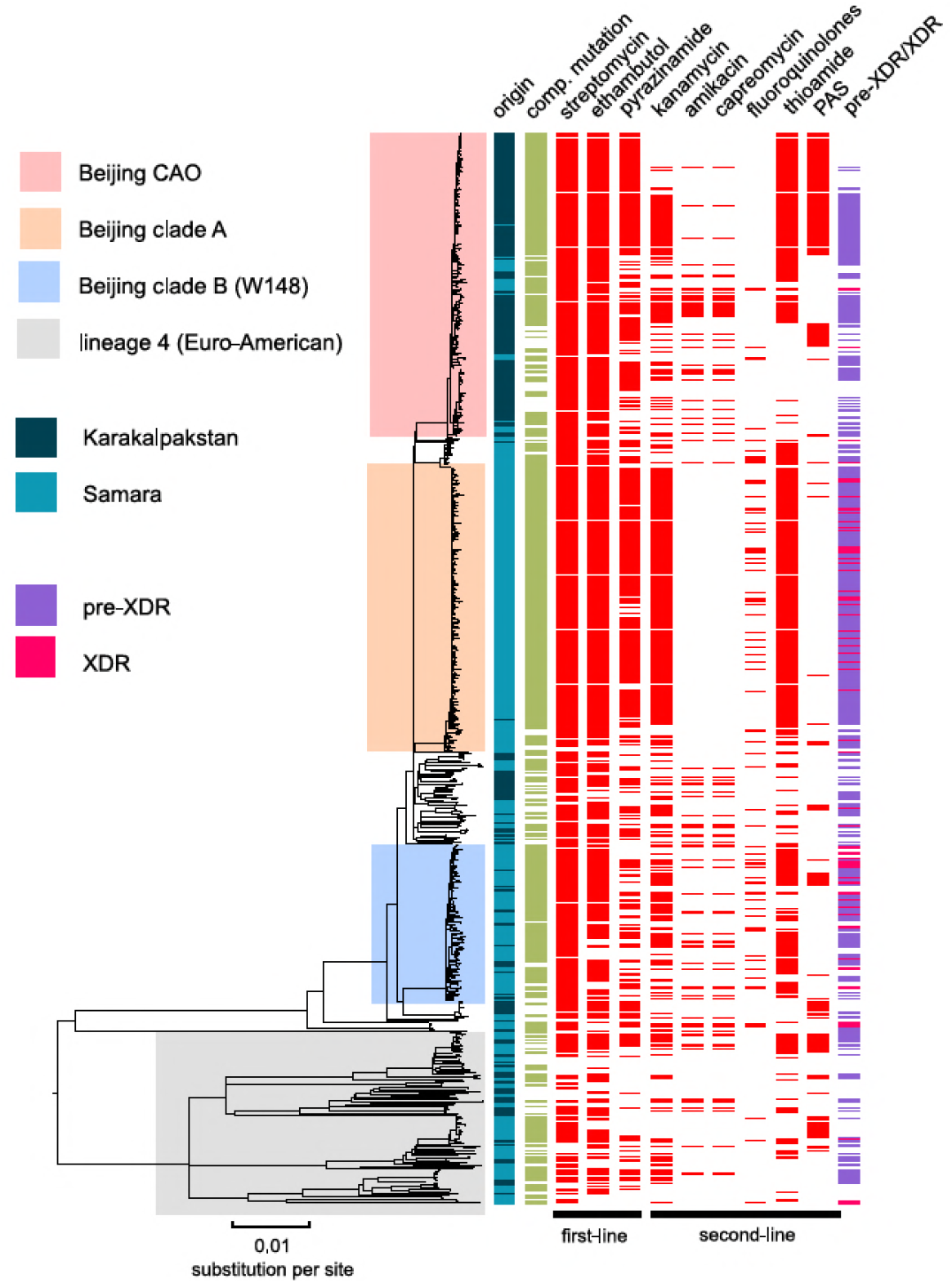
MDR-MTBC phylogeny and resistance mutations of strains from Samara (Russia) and Karakalpakstan (Uzbekistan) Maximum likelihood tree (with 1,000 resamples, GTR nucleotide substitution model) based on 12,567 variable positions (SNPs) among 705 MDR-MTBC isolates from **Karakalpakstan and Samara**. Any resistance associated mutations (see methods) for individual antibiotics are depicted with red bars for each strain. The presence of any putative compensatory mutation in the RNA polymerase genes *rpoA, rpoB, rpoC* is depicted with green bars and country of origin and a genotypic pre-XDR and XDR strain classification is color coded. PAS = para-aminosalycylic acid.

The three Beijing clades (A, B, and CAO) in Samara and Karakalpakstan had more drug resistance conferring mutations (in addition to isoniazid and rifampicin resistance) with means of 5.0 (SEM 0.07), 4.2 (SEM 0.18), and 4.7 (SEM 0.11), respectively (Appendix Figure S5), than compared to only 3.6 (SEM 0.20) additional genotypic drug resistances (*P* < 0.0001, *P* = 0.0143, *P* < 0.0001) for other Beijing strains in both settings. Strains belonging to other MTBC genotypes (mainly lineage 4 subgroups) were found with a mean of 2.6 (SEM 0.20) additional drug resistance mediating mutations, lower than any Beijing-associated group (*P* ≤ 0.0009) (Appendix Figure S5).

Similar to Karakalpakstan, MDR-MTBC strains from Samara with compensatory mutations also accumulated more resistance-associated mutations (4.57 vs 2.30 mutations per genome; two-sample t-test *P*<2.2×10^−16^) and had higher transmission indexes (50.32 vs 0.46; Welch two-sample t-test *P*<2.2×10^−16^) compared to strains lacking compensatory mutations (Appendix Figure S6).

The impact of resistance conferring and compensatory mutations on the transmission success score in Beijing-A clade from Samara (Appendix Figure S7) was strikingly similar to the one observed in CAO strains from Karakalpakstan. The presence of compensatory mutations, but not the accumulation of resistance mutations, was significantly and independently associated with network size in clusters originating in the 1980s and 1990s, with a maximum influence found in clusters starting in the late 1990s. Critically, the high proportions of strains detected in both settings with pre-XDR and XDR resistance profiles among the three major Beijing clades (clade A, 96% clade B, 62% clade CAO, 50% Appendix Table S4, Figure 6) reveal the low proportion of patients that are or would be eligible to receive the newly WHO endorsed short MDR-TB regimen. As per definition of the WHO exclusion criteria, e.g. any confirmed or suspected resistance to one drug (except isoniazid) in the short regimen, only 0.5% (1/191 in Karakalpakstan) and 2.7% (8/300 in Samara) of the patients infected with either a Beijing clade A, B or CAO strain would benefit from a shortened MDR-TB therapy (additional data).

**Fig. 6:**
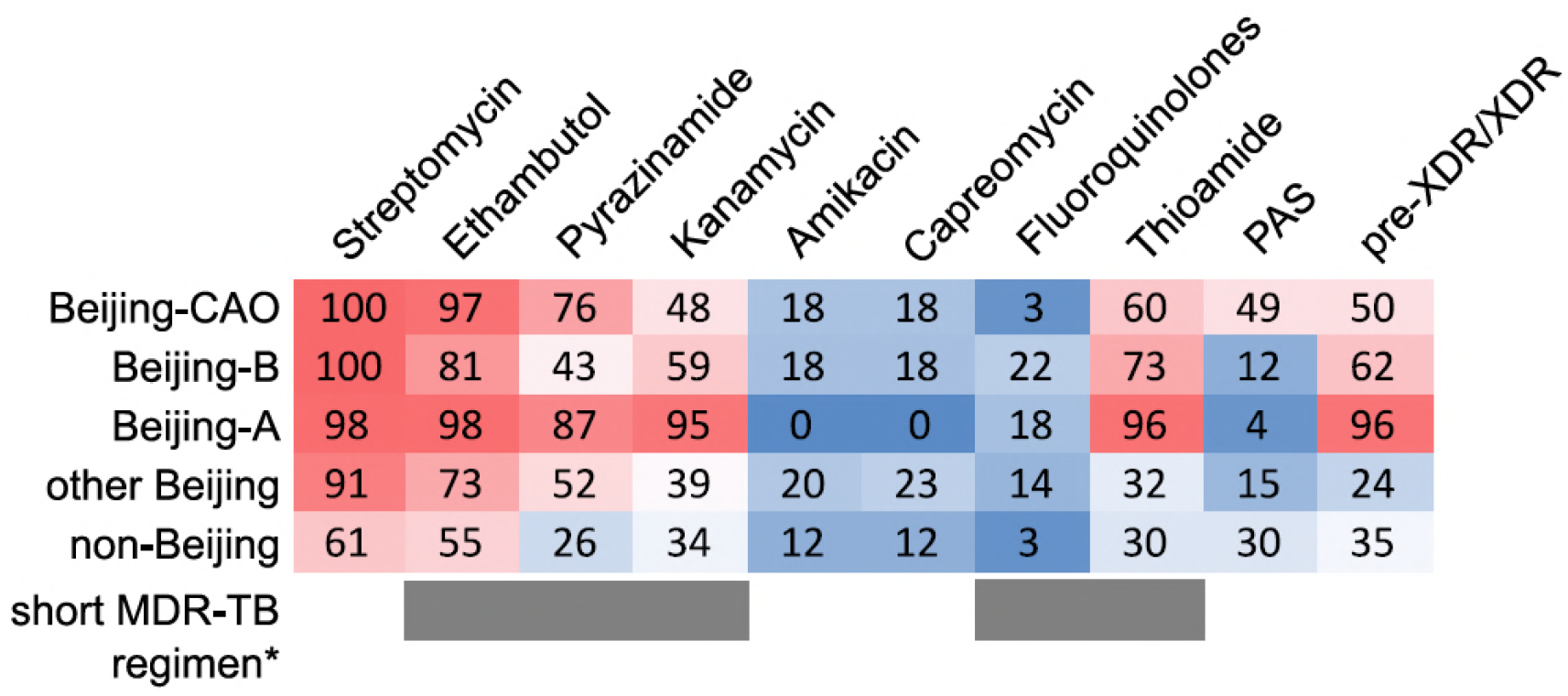
Percentage of drug resistance among 705 MDR-MTBC strains from Samara (Russia) and Karakalpakstan (Uzbekistan) MDR-MTBC strains stratified to three Beijing sub-groups, other Beijing strains and non-Beijing strains. Proportions of strains with identified molecular drug resistance mutations (see additional data) which mediate resistance to multiple first- and second-line anti-TB drugs. Values are rounded. Drugs used in the WHO endorsed standardized short MDR-TB regimen marked with grey boxes. *The short MDR-TB regimen further includes high-dose isoniazid treatment, and clofazimine. In that regard, we identified 622/705 (85.4%) of the MDR-MTBC strains with the well-known high-level isoniazid resistance mediating mutation *katG* S315T (additional data), for clofazimine resistance mediating mutations are not well described.

## Discussion

Using WGS combined with Bayesian and phylogenetic analyses, we reveal the evolutionary history and recent clonal expansion of the dominatant MDR/pre-XDR MTBC clade in Karakalpakstan, Uzbekistan, termed the Central Asian outbreak (CAO). Strikingly, CAO-strains were also found also in Samara, Russia, and vice versa strains belonging to the second largest clade in Samara (Beijing clade B, i.e. European/Russian W148 (13, 22)) were identified in Karakalpakstan, suggesting that the MDR-TB epidemic in this world region is driven by few outbreak clades. During the three last decades, these strains gradually accumulated resistance to multiple anti-TB drugs that largely escaped phenotypic and molecular diagnostics, and reduce treatment options to a restricted set of drugs that often cause severe side effects. In addition, our results suggest that compensatory mutations (in RNA-polymerase subunit coding genes) that are proposed to ameliorate growth deficits in rifampicin resistant strains in vitro are also crucial in a global epidemiological context allowing MDR and pre-XDR strains to form and maintain large transmission networks. The predominance of these strain networks, seen in two distant geographic regions of the former Soviet Union clearly limit the use of standardized MDR-TB therapies, e.g. the newly WHO endorsed short MDR-TB regimen, in these settings.

Temporal reconstruction of the resistance mutation acquisition and of changes in bacterial population sizes over three decades demonstrates that MDR outbreak strains already became resistant to both first- and second-line drugs in the 1980s. Fully first-line resistant strains massively expanded in the 1990s, a period that shortly preceded or immediately followed the end of the Soviet Union, years before the implementation of DOTS and programmatic second-line MDR-TB treatment. This is in line with the known rise in TB incidence that accompanied the economic breakdown in Russia during the 1990’s (23).

From a bacterial genetic point of view, our data show that particular MDR and pre-XDR strain subgroups are highly transmissible despite accumulation of multiple resistance mutations. The acquisition of compensatory mutations after introduction of low fitness cost resistance mutations (e.g. *katG* S315T (10), *rpoB* S450L (8), *rpsL* K43R (24)) seems the critical stage allowing for higher transmission rates. Multiple regression analyses further strengthened this hypothesis by demonstrating that the presence of fitness compensating variants was positively associated with transmission success in different settings and outbreak clades, independently of the accumulation of resistance mutations. Compensatory evolution thus appears to play a central role in driving large MDR-TB epidemics such as that seen with the Beijing CAO-clade.

A particular concern is the high prevalence of mutations conferring resistance to second-line drugs currently included in treatment regimens, among the dominant MDR-MTBC strains. Their detected emergence in a period preceding DOTS implementation, e.g. in Karakalpakstan, can be explained by past, largely empirical treatment decisions or self-medication. For instance, high frequencies of mutations in the *ribD* promoter region, and *folC* among Beijing-CAO strains, associated with para-aminosalicylic acid resistance (25, 26), are a likely consequence of the use of para-aminosalicylic acid in failing treatment regimens in the late 1970s to the early 1980s in the Soviet Union (27–29). Likewise, the frequent independent emergence of mutations in the *eis* promoter and of rare variants in the upstream region of *whiB7*, both linked to resistance to aminoglycosides (mainly streptomycin and kanamycin) (30, 31), probably reflects self-administration of kanamycin that was available in local pharmacies.

The pre-existence of fully first-line resistant strain populations (e.g. CAO-Beijing in Karakalpakstan) likely contributed to the poor treatment outcomes observed among MDR-TB patients following the implementation of first-line DOTS treatment in 1998 (16). This period coincides with a detected CAO population size increase, likely reflecting the absence of drug susceptibility testing and therefore appropriate second-line treatment during extended hospitalization at the time, resulting in prolonged infectiousness of TB-patients and further spread of these strains.

The frequencies of fluoroquinolone resistance, mediated by *gyrA* and *gyrB* mutations, remained low among the Karakalpakstan MDR-MTBC strains, which is consistent with the notion that such drugs were rarely used for treating TB in former Soviet Union countries (see discussion (13), (27–29)). This observation explains the generally favorable MDR-TB treatment outcomes observed with the use of individualized second-line regimens, including a fluoroquinolone, in the latter MDR-TB treatment program in the Karakalpakstan patient population (14, 32). However, fluoroquinolone resistance, representing the last step towards XDR-TB, is already emerging as reported for strains in Beijing clade A and B (13).

In conclusion, the (pre-) existence and wide geographic dissemination of highly resistant and highly transmissible strain populations most likely contributes to increasing M/XDR-TB incidence rates despite scaling up of the MDR-TB programs in some Eastern European and Russian regions (15, 23, 33). Importantly, from the large spectrum of resistance detected among dominating strains in this study, it can be predicted that standardized therapies, including the newly WHO endorsed short MDR-TB regimen in Uzbekistan, are/will be largely ineffective for many patients in Samara and Karakalpakstan, and likely elsewhere in Eurasia. In order to successfully control the worldwide MDR-TB epidemics, universal access to rapid and comprehensive drug susceptibility testing, best supported by more advanced technologies, will be crucial for guiding individualized treatment with existing and new/repurposed TB drugs and to maximize chances of cure and prevention of further resistance acquisition.

## Methods

### Study population, Karakalpakstan (Uzbekistan)

A total of 277 MDR-MTBC strains derived from two separate cohorts were sequenced. The first cohort comprised 86% (49/57) of MDR-MTBC strains from a cross-sectional drug resistance survey conducted in four districts in Karakalpakstan, Uzbekistan between 2001-2002 (16). An additional 228 strains were obtained from TB-patients enrolled for second-line treatment in the MDR-TB treatment program from 2003 to 2006. These strains represented 76% (228/300) of all MDR-TB cases diagnosed over the period. While the MDR-TB treatment program covered two of the four districts included in the initial drug resistance survey, the majority of strains from both cohorts, 69% and 64% respectively, were obtained from patients residing in the same main city of Nukus (Appendix Table S1).

### Study population, Samara (Russia)

To set the MDR-MTBC strains from Karakalpakstan into a broader geographical perspective, raw sequencing data of 428 MDR-MTBC strains from a published cross-sectional prospective study in Samara, Russia from 2008-2010 (13) were processed as described below and included into a composite MDR-MTBC dataset.

### Drug susceptibility testing

Drug susceptibility testing (DST) was performed for five first-line drugs (isoniazid, rifampicin, ethambutol, streptomycin, pyrazinamide), and three second-line drugs (ofloxacin, capreomycin and prothionamide) for cohort 1, and six second-line drugs for cohort 2 (capreomycin, amikacin, ofloxacin, ethionamide, para-aminosalicylic acid and cycloserine) by the reference laboratory in Borstel, Germany as described previously (17).

### Whole genome sequencing

Whole genome sequencing (WGS) was performed with Illumina Technology (MiSeq and HiSeq 2500) using Nextera XT library preparation kits as instructed by the manufacturer (Illumina, San Diego, CA, USA). Fastq files (raw sequencing data) were submitted to the European nucleotide archive (see additional data for accession numbers). Obtained reads were mapped to the *M. tuberculosis* H37Rv reference genome (GenBank ID: NC_000962.3) with BWA (18). Alignments were refined with GATK (19) and Samtools (20) toolkits with regard to base quality re-calibration and alignment corrections for possible PCR artefact. We considered variants that were covered by a minimum of 4 reads in both forward and reverse orientation, 4 reads calling the allele with at least a phred score of 20, and 75% allele frequency. In the combined datasets, we allowed a maximum of 5% of all samples to fail the above mentioned threshold criteria in individual genome positions to compensate for coverage fluctuations in certain genome regions; in these cases, the majority allele was considered. Regions annotated as ‘repetitive’ elements (e.g. PPE and PE-PGRS gene families), insertions and deletions (InDels), and consecutive variants in a 12 bp window (putative artefacts flanking InDels) were excluded. Additionally, 28 genes associated with drug resistance and bacterial fitness (see additional data) were excluded for phylogenetic reconstructions. The remaining single nucleotide polymorphisms (SNPs) were considered as valid and used for concatenated sequence alignments. Further detailed methods of the phylogenetic reconstruction, molecular resistance prediction, strain-to-strain genetic distance, and Bayesian models are given as Supplementary Methods.

### Transmission index

Based on the distance matrix (SNP distances), we further determined for every isolate the number of isolates that were in a range of 10 SNPs or less (in the following referred to as “transmission index”). This 10 SNP-threshold was used to infer the number of recently linked cases, as considered within a 10-year time period, based on previous convergent estimates of MTBC genome evolution rate of ≈ 0.5 SNPs/genome/year in inter-human transmission chains and in macaque infection models (21–24).

### Genotypic drug resistance prediction

Mutations (small deletions and SNPs) in 34 resistance associated target regions (comprising 28 genes) were considered for a molecular resistance prediction to 13 first- and second-line drugs (additional data). Mutations in genes coding for the RNA-Polymerase subunits *rpoA, rpoB* (excluding resistance mediating mutations), and *rpoC* were reported as putative fitness compensating (e.g. in vitro growth enhancing) variants for rifampicin resistant strains. A detailed overview of all mutations considered as genotypic resistance marker is given as additional data. Mutations that were not clearly linked to phenotypic drug resistance were reported as genotypic non wild type and were not considered as genotypic resistance markers. When no mutation (or synonymous, silent mutations) was detected in any of the defined drug relevant target regions the isolate was considered to be phenotypically susceptible.

### Phylogenetic inference (maximum likelihood)

We used jModelTest v2.1 and Akaike and Bayesian Information Criterion (AIC and BIC) to find an appropriate substitution model for phylogenetic reconstructions based on the concatenated sequence alignments (Appendix Table S5). Maximum likelihood trees were calculated with FastTree 2.1.9 (double precision for short branch lengths) (25) using a general time reversible (GTR) nucleotide substitution model (best model according to AIC and second best model according to BIC), 1,000 resamplings and Gamma20 likelihood optimization to account for evolutionary rate heterogeneity among sites. The consensus tree was rooted with the “midpoint root” option in FigTree and nodes were arranged in increasing order. Polymorphisms considered as drug resistance marker (see above) and putative compensatory variants were analyzed individually and mapped on the phylogenetic tree to define resistance patterns of identified phylogenetic subgroups.

### Molecular clock model

In order to compute a time scaled phylogeny and employ the Bayesian skyline model (see below) for the identified Central Asian outbreak (CAO) clade we sought to define an appropriate molecular clock model (strict versus relaxed clock) and a mutation rate estimate. Due to the restricted sampling timeframe of the Karakalpakstan dataset (2001-2006) we extended the dataset for the model selection process with CAO strains from Samara (2008-2010) and ‘historical’ CAO strains isolated from MDR-TB patients in Germany (1995-2000) thus allowing for a more confident mutation rate estimate. The strength of the temporal signal in the combined dataset, assessed by the correlation of sampling year and root-to-tip distance, was investigated with TempEst v1.5 (26). Regression analysis was based on residual mean squares, using a rooted ML tree (PhyML, GTR substitution model, 100 bootstraps), R-square and adjusted *P*-value are reported. For the comparison of different Bayesian phylogenetic models we used path sampling with an alpha of 0.3, 50% burn-in and 15 million iterations (resulting in mean ESS values >100), marginal likelihood estimates were calculated with BEAST v2.4.2 (27), and Δ marginal L estimates are reported relative to the best model.

First, we employed a strict molecular clock fixed to 1 × 10^−7^ substitutions per site per year as reported previously (21–23) without tip dating, a strict molecular clock with tip dating and a relaxed molecular clock with tip dating. BEAST templates were created with BEAUti v2 applying a coalescent constant size demographic model, a GTR nucleotide substitution model, a chain length of 300 million (10% burn-in) and sampling of 5,000 traces/trees.

Second, we ran different demographic models (i.e. coalescent constant size, exponential, and Bayesian skyline) under a relaxed molecular clock using tip dates and the same parameters for the site model and Markov-Chain-Monte-Carlo (MCMC) as described above. Inspection of BEAST log files with Tracer v1.6 showed an adequate mixing of the Markov chains and all parameters were observed with an effective sample size (ESS) in the hundreds, suggesting an adequate number of effectively independent draws from the posterior sample and thus sufficient statistical support.

### Bayesian Skyline Plot

Changes of the effective population size of the CAO clade in Karakalpakstan over the last four decades were calculated with a Bayesian skyline plot using BEAST v2.4.2 (27) using a tip date approach with a strict molecular clock model of 0.94 × 10^−7^ substitutions per site per year (best model according to path sampling results, see above), and a GTR nucleotide substitution model. We further used a random starting tree, a chain length of 300 million (10% burn-in) and collected 5,000 traces/trees. Again adequate mixing of the Markov chains and ESS values in the hundreds were observed. A maximum clade credibility genealogy was calculated with TreeAnnotator v2.

### Impact of resistance-conferring and compensatory mutations on transmission success

We used multiple linear regression to examine the respective contributions of antimicrobial resistance and putative fitness cost-compensating mutations to the transmission success of tuberculosis. To take transmission duration into account, we computed, for each isolate and each period length *T* in years (from 1 to 40y before sampling), a transmission success score defined as the number of isolates distant of less than *T* SNPs, divided by *T*. This approach relied on the following rationale: based on MTBC evolution rate of 0.5 mutation per genome per year, the relation between evolution time and SNP divergence is such that a cluster with at most *N* SNPs of difference is expected to have evolved for approximately *N* years. Thus, transmission success score over *T* years could be interpreted as the size of the transmission network divided by its evolution time, hence as the average yearly increase of the network size. For each period *T*, the transmission success score was regressed on the number of resistance mutations and on the presence of putative compensatory mutations. The regression coefficients with 95% confidence intervals were computed and plotted against *T* to identify maxima, that is, time periods when the transmission success was maximally influenced by either resistance-conferring or –compensating mutations. These analyses were conducted independently on outbreak strains of the Beijing-CAO clade in the Karakalpakstan cohort and of the Beijing-A clade in the Samara cohort.

### Statistical analyses

Differences between cohorts and numbers of sampled isolates per year category were performed using Chi-squared analysis (mid-P exact) or Fisher’s exact test, while comparison of median age was performed using the Mann-Whitney test. *P*-values for pairwise comparisons of subgroups regarding pairwise genetic distances, number of resistant DST results and number of resistance related mutations were calculated with an unpaired t-test (Welch correction) or a t-test according to the result of the variances comparison using a F-test. Boxplot, bubble plots and density plots have been performed on R.

## Acknowledgments

We thank: I. Razio, P. Vock, T. Ubben and J. Zallet from Borstel, Germany for technical assistance; the national and expatriate staff of Médecins Sans Frontières, Karakalpakstan; Dr. Atadjan and Dr. K. Khamraev from the Ministry of Health (Karakalpakstan) for their support. Parts of this work have been supported by the European Union TB-PAN-NET (FP7-223681) project and the German Center for Infection Research. The funders had no role in study design, data collection and analysis, decision to publish, or preparation of the manuscript. Raw sequence data (fastq files) have been deposited at the European Nucleotide Archive (ENA) under the project number (pending).

## Appendix Figures and Tables

**Appendix Figure S1:**
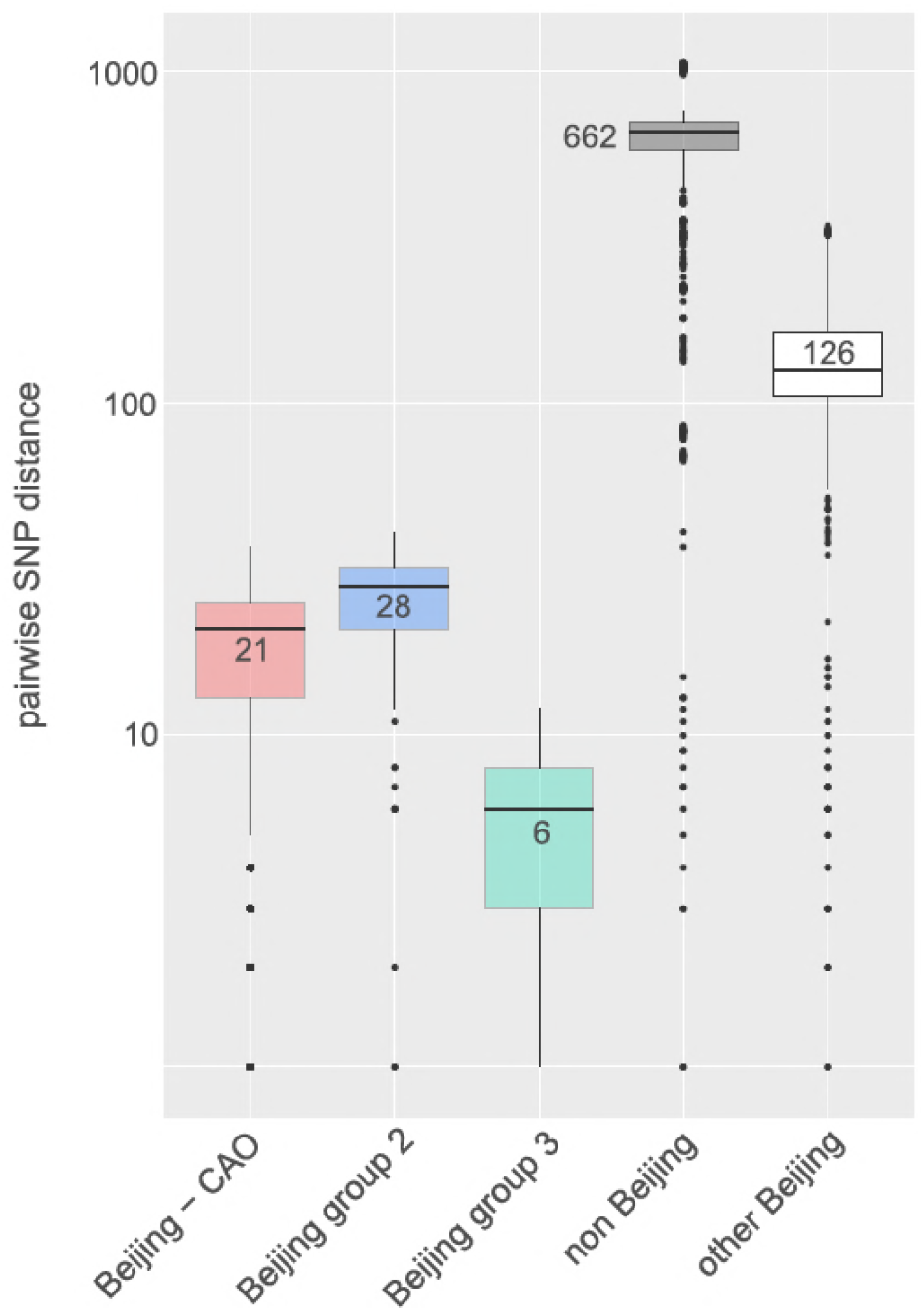
Box-Plot showing pairwise SNP distances among identified Beijing genotype subgroups in comparison to non-Beijing strains from **Karakalpakstan**, Uzbekistan. Box represents inter quartile range, whiskers represent 95% of the data, outliers shown as black dots; solid black line represents the median.

**Appendix Figure S2:**
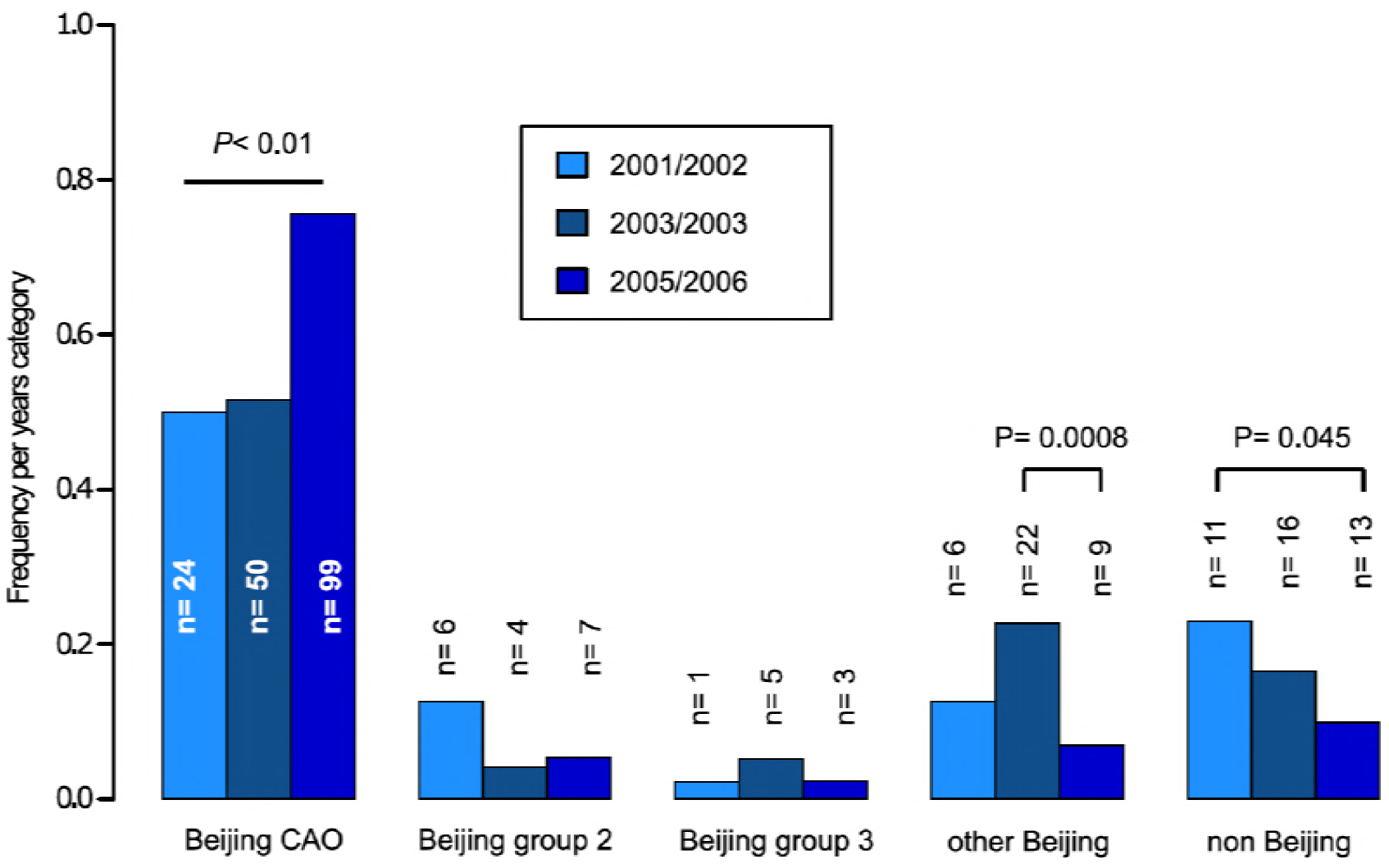
Proportions of different genome-based subgroups in **Karakalpakstan**, Uzbekistan stratified to the years 2001/02, 2003/04, 2005/06. P-values for pairwise comparisons within phylogenetic groups were calculated with Fisher exact test (two-sided). Beijing CAO 2001/2002 vs 2005/2006 *P*=0.0018, Beijing CAO 2003/2004 vs 2005/2006 *P*=0.0002.

**Appendix Figure S3:**
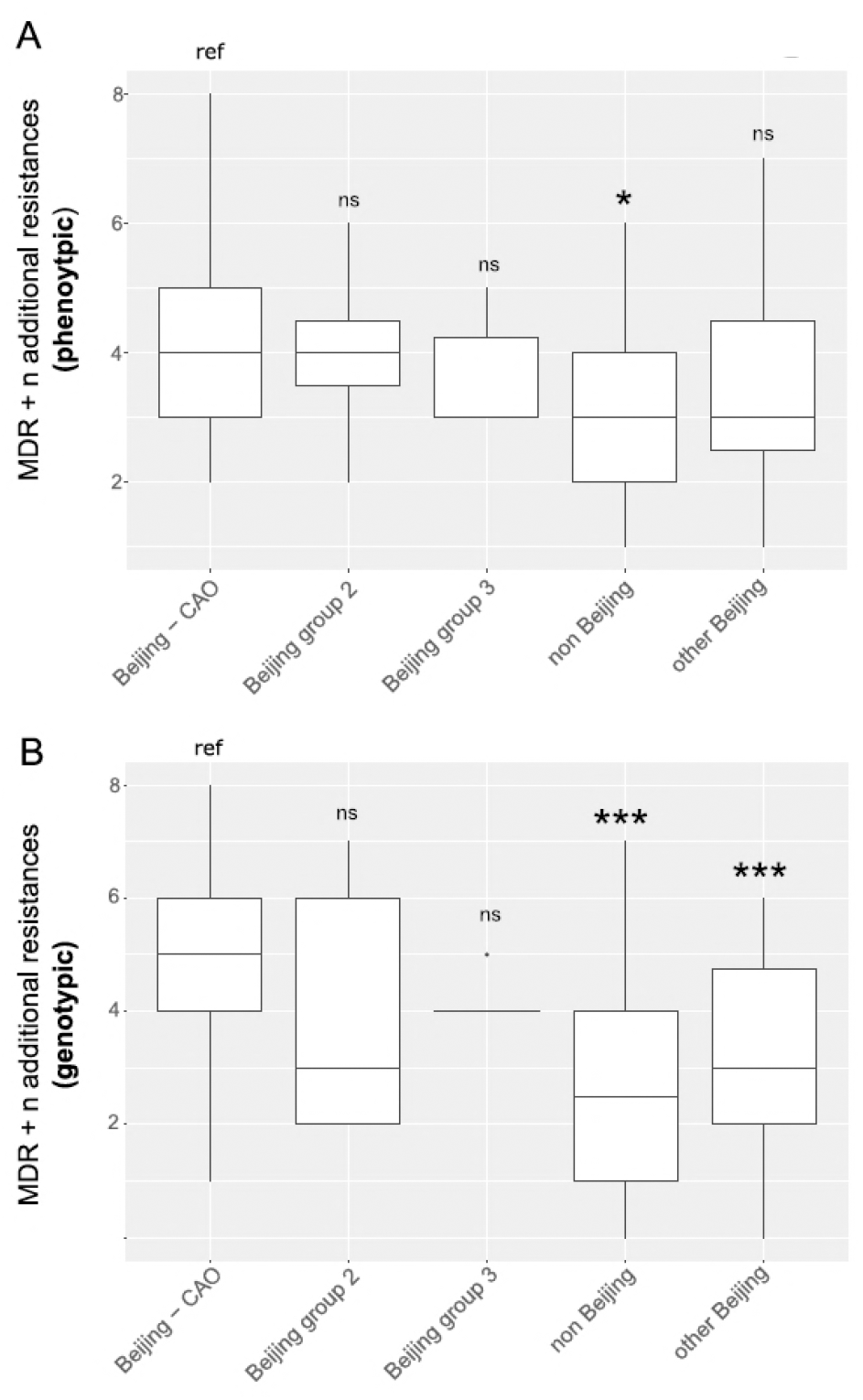
Box-Plot showing median number of (A) phenotypic and (B) genotypic drug resistances (in addition to the MDR classification, i.e. isoniazid and rifampicin resistance) of all strains **from Karakalpakstan**. Box represents inter quartile range, whiskers represent 95% of the data, outliers shown as black dots; solid black line represents the median. Beijing CAO strains exhibit more phenotypic drug resistances compared to non-Beijing strains (*P*=0.0079) and more genotypic drug resistances compared to other Beijing strains (*P*<0.0001), and non-Beijing strains (*P*<0.0001). P-values for pairwise comparison with reference group calculated with unpaired t-test (two-tailed, Welch’s correction).

**Appendix Figure S4:**
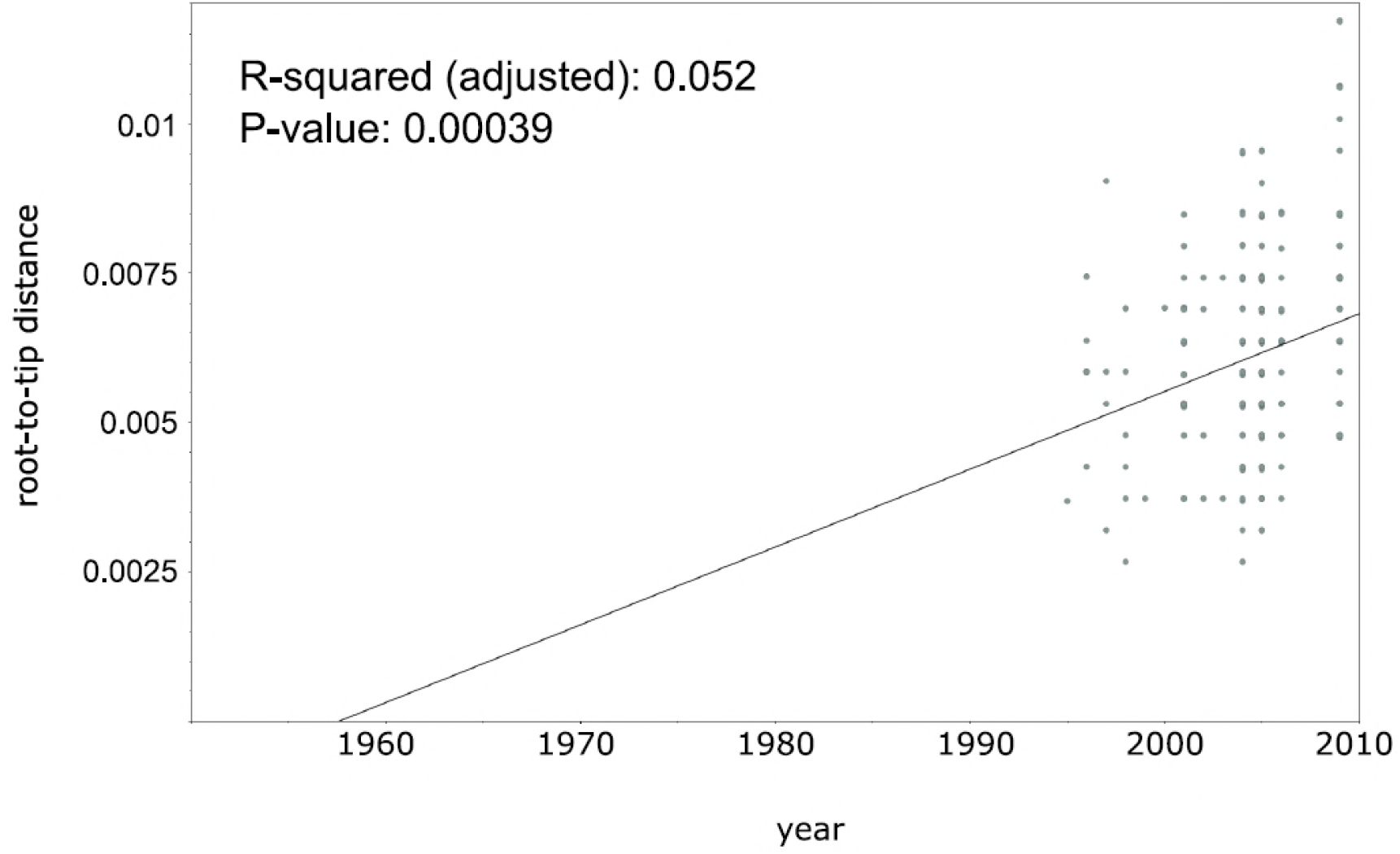
Linear regression analysis showing correlation between root-to-tip distance and sampling years of an extended collection of 220 Beijing CAO datasets covering the period 1995 to 2009.

**Appendix Figure S5:**
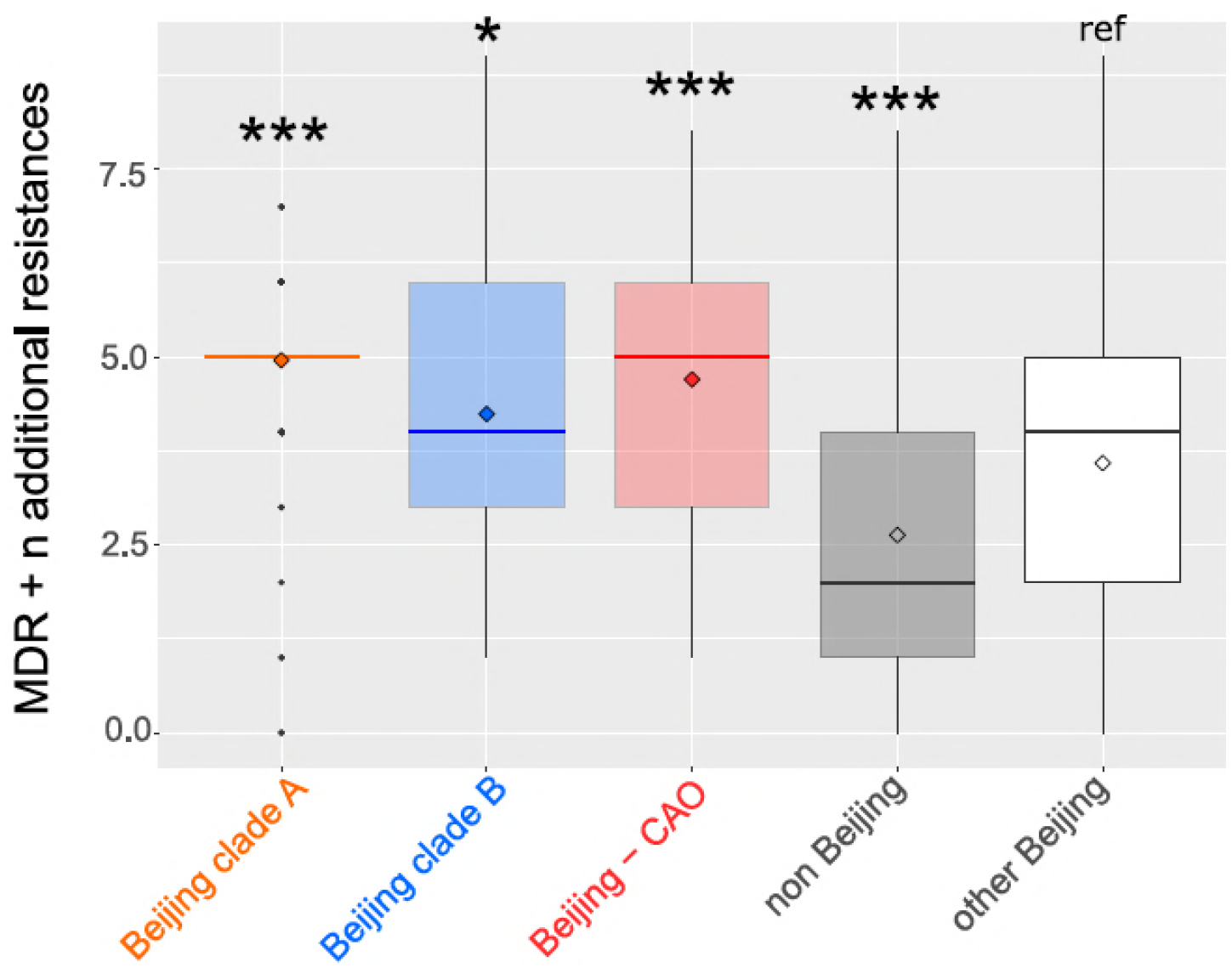
Number of drug resistance mutations among different MDR-MTBC groups from **Samara** (n=428) **and Karakapakstan** (n=277). Box-Plot with mean (diamond) and median (horizontal line) number of genotypic drug resistances (see methods) to additional anti-TB drugs (beyond MDR defining rifampicin and isoniazid resistance). Box represents inter quartile range, whiskers represent 95% of the data, outliers shown as black dots. *P*-values for three major Beijing outbreak clades (A, B and CAO), and non-Beijing strains (mainly lineage 4 isolates) were calculated with unpaired t-tests with Welch correction compared to the group ‘other Beijing’ strains. Color codes according to Fig. 5. P-values for pairwise comparison with reference group calculated with unpaired t-test (two-tailed, Welch’s correction). Clade A (*P*≤0.0001), Clade B (*P*=0.0143), CAO (*P*≤0.0001), and non-Beijing (*P*=0.0009).

**Appendix Figure S6:**
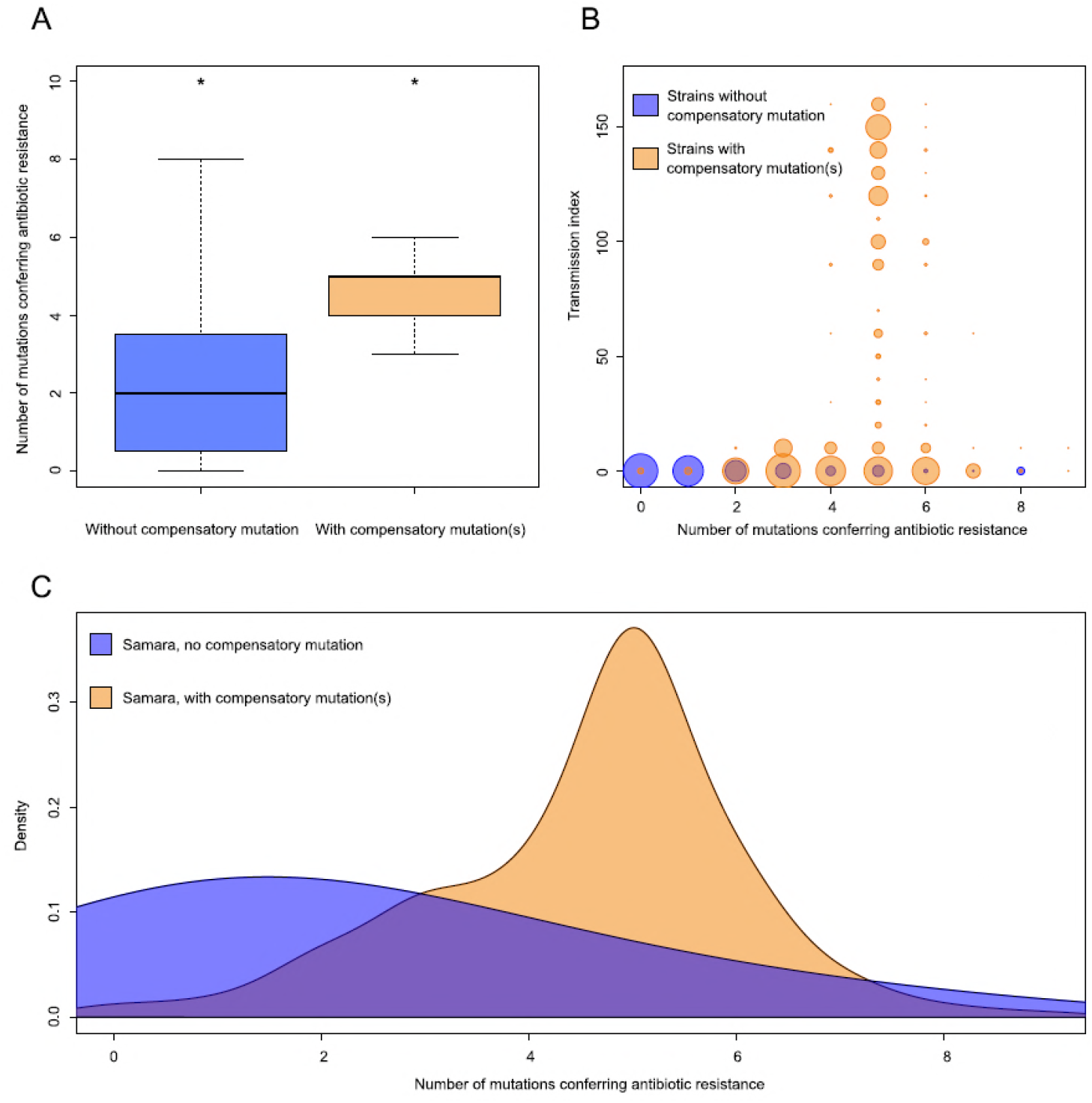
Comparisons between strains carrying compensatory mutations (in orange) and strains with no-compensatory mutations (in blue), from the **Samara dataset**. A) Boxplot showing number of resistance mutations for the two categories (without or with compensatory mutations). The two categories were significantly different (two-sample t-test *P*<2.2×10^−16^). B) Bubble plots showing the transmission index (number of strains differing by less than 10 SNPs) as a function of antibiotic resistance related mutations. Bubble sizes are function of the number of strains. C) Density plot of the number of resistance-conferring mutations for strains carrying compensatory mutations (orange) and strains that don’t carry compensatory mutation (blue) from Samara dataset. Proportions are adjusted by using Gaussian smoothing kernels.

**Appendix Figure S7:**
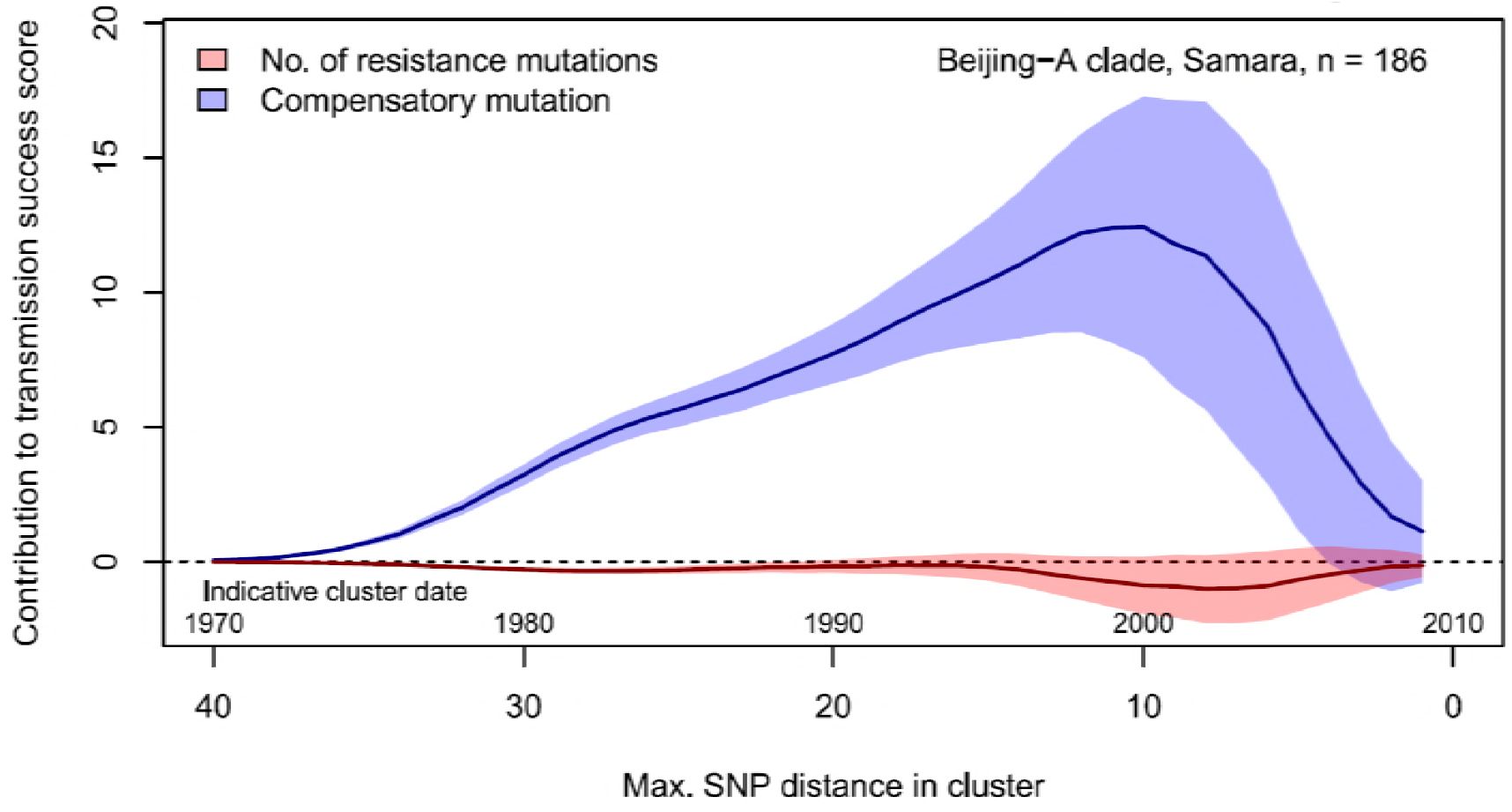
Contributions of resistance-conferring and compensatory mutations to the transmission success of *M. tuberculosis* of the Beijing-A clade from **Samara**, Russia. Shown are the coefficients and 95% confidence bands of multiple linear regression of the transmission success score, defined as the size of clusters diverging by at most *N* SNPs and divided by *N* or, equivalently, the size of clusters that evolved over *N* years divided by *N*. Compensatory mutations were independently associated with transmission success, with a maximum association strength found for transmission clusters beginning around 1999.

**Appendix Table S1:**
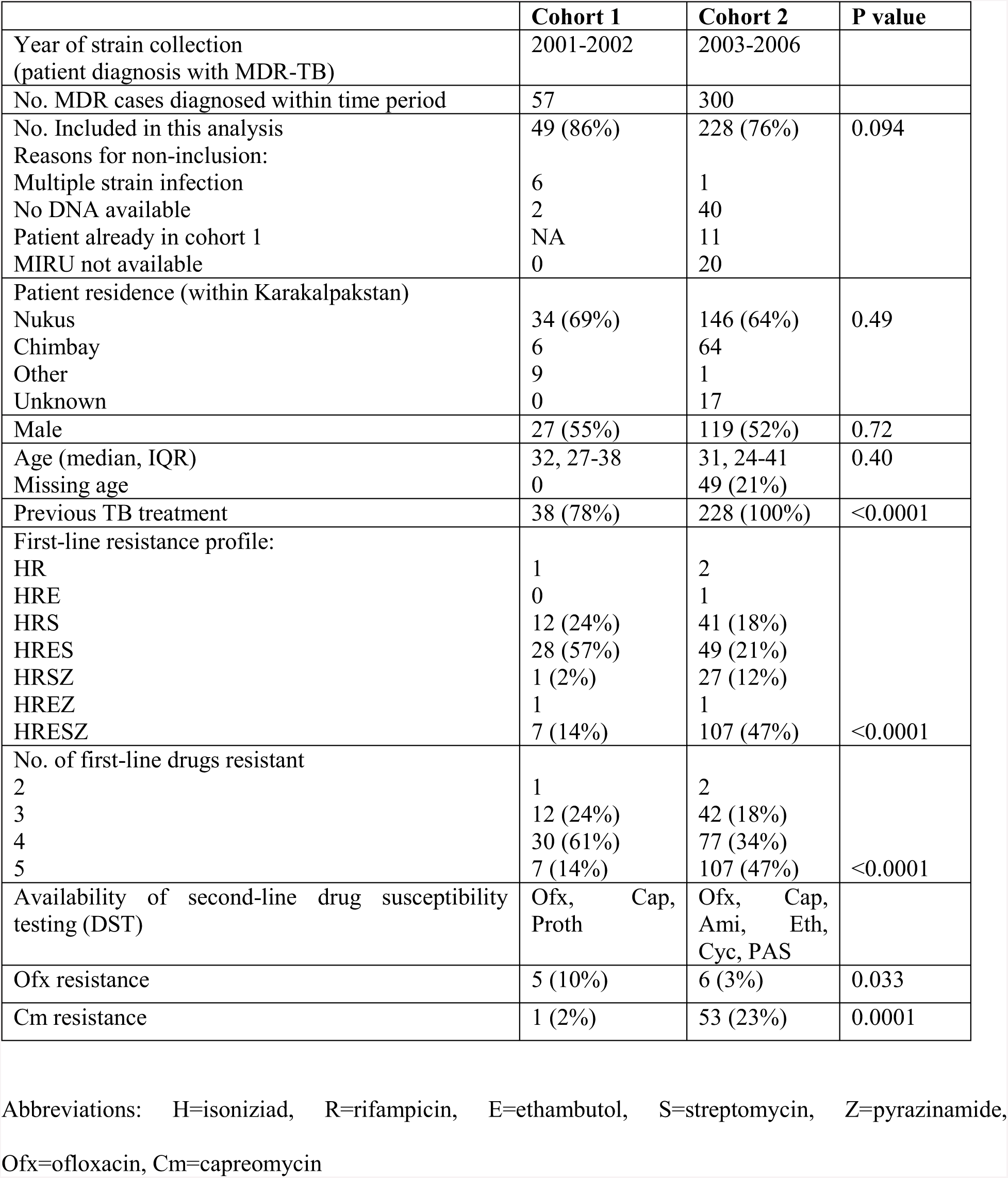
Main characteristics of patients from cohorts 1 and 2 in **Karakalpakstan,** Uzbekistan.

**Appendix Table S2:** Path sampling results and model selection based on Δ marginal L estimates (relative to best model, given in bold fonts) considering 75 path sampling steps and chain lengths of 15 million analysing Beast runs of a combined dataset of Central Asian outbreak (CAO) isolates originated from Germany (1995-2000), **Karakalpakstan** (2001-2006), **and Samara** (2008-2010).

| Subst. model | Clock model | Demographic model | Marginal L estimate | Mean ESS | $\Delta$ marginal L estimate | Subst rate x $10^{-7}$ (95%HPD) |
| --- | --- | --- | --- | --- | --- | --- |
| GTR | Strict (no tip dating) | Coalescent constant size | -10131.67 | 214.3 | 32.21 | 1.0 (fixed) |
| **GTR** | **Strict (tip dating)** | **Coalescent constant size** | **-10099.46** | **153.7** | **ref** | **1.0 (fixed)** |
| GTR | Relaxed, lognormal | Coalescent constant size | -10117.21 | 123.9 | 17.75 | 0.96 (0.65-1.24) |
| *Strict clock with tip dating selected as best model* |  |  |  |  |  |  |
| GTR | Relaxed, lognormal | Coalescent constant size | -10117.21 | 123.9 | 78.28 | 0.96 (0.65-1.24) |
| GTR | Relaxed, lognormal | Exponential | -10044.41 | 189.5 | 5.48 | 0.88 (0.58-1.21) |
| **GTR** | **Relaxed, lognormal** | **Bayesian skyline** | **-10038.93** | **98.6** | **ref** | **0.94 (0.72-1.15)** |
| *Mutation rate of 0.94 used for a Bayesian skyline model with a strict molecular clock for CAO strains from Karakalpakstan only* |  |  |  |  |  |  |

Abbreviations: HPD=Highest posterior density interval, GTR=general time reversible

**Appendix Table S3:** Mutations in *rpoB, rpoA* and *rpoC* associated with a putative compensatory effect in rifampicin resistant MTBC strains. Data from 277 MDR-MTBC strains from **Karakalpakstan**, Uzbekistan, stratified to the particularly successful variant termed Central Asian outbreak (CAO) and other Beijing strains. Pairwise differences between the two groups calculated with Fisher exact test; two-tailed *P*-values are reported.

|  | Beijing CAO<br>(n=173) | Other Beijing<br>(n=64) | *P*-value | All<br>(n=277) |
| --- | --- | --- | --- | --- |
| *rpoB* mutations outside RRDR, excluding codon 170,400,491 variants | 25 (14.5%) | 12 (18.8%) |  | 43 (15.5%) |
| wild type | 147 (85.0%) | 52 (81.3%) | 0.43 | 234 (84.5%) |
| *rpoC* variants | 95 (54.9%) | 18 (28.1%) |  | 126 (45.5%) |
| wild type | 78 (45.1%) | 46 (71.2%) | 0.0002 | 151 (54.5%) |
| *rpoA* variants | 5 (2.9%) | 2 (3.1%) |  | 7 (2.5%) |
| wild type | 168 (97.1%) | 62 (96.9%) | 1.00 | 270 (97.5%) |

Abbreviations: CAO=Central Asian outbreak, RRDR=rifampicin resistance determining region

**Appendix Table S4:** Proportions of genotypic drug resistance rates for different anti-TB drugs (beyond isoniazid and rifampicin resistance) and pre-XDR/XDR-TB classification among 705 MDR-MTBC clinical isolates from **Samara** (n=428) **and Karakalpakstan** (n=277), stratified to three identified major phylogenetic groups within the Beijing genotype/lineage and to other Beijing strains, and to non-Beijing strains (mainly lineage 4, Euro-American).

| group | S | E | Z | Km | Am | Cm | Fq | Thio | PAS | Pre-XDR<br>XDR |
| --- | --- | --- | --- | --- | --- | --- | --- | --- | --- | --- |
| Beijing CAO<br>(n=201) | 201/201<br>100.0% | 195/201<br>97.0% | 152/201<br>75.6% | 97/201<br>48.3% | 37/201<br>18.4% | 37/201<br>18.4% | 6/201<br>3.0% | 121/201<br>60.2% | 99/201<br>49.3% | 100/201<br>49.8% |
| Beijing clade<br>B (W148)<br>(n=103) | 103/103<br>100.0% | 83/103<br>80.6% | 44/103<br>42.7% | 61/103<br>59.2% | 18/103<br>17.5% | 18/103<br>17.5% | 23/103<br>22.3% | 75/103<br>72.8% | 12/103<br>11.7% | 64/103<br>62.1% |
| Beijing clade<br>A (n=187) | 184/187<br>98.4% | 183/187<br>97.9% | 163/187<br>87.2% | 177/187<br>94.7% | 0/187<br>0.0% | 0/187<br>0.0% | 33/187<br>17.6% | 180/187<br>96.3% | 7/187<br>3.7% | 179/187<br>95.7% |
| Other Beijing<br>(n=100) | 91/100<br>91.0% | 73/100<br>73.0% | 52/100<br>52.0% | 39/100<br>39.0% | 20/100<br>20.0% | 23/100<br>23.0% | 14/100<br>14.0% | 32/100<br>32.0% | 15/100<br>15.0% | 45/187<br>24.1% |
| Non-Beijing<br>(n=114) | 69/114<br>60.5% | 63/114<br>55.3% | 30/114<br>26.3% | 39/114<br>34.2% | 14/114<br>12.3% | 14/114<br>12.3% | 3/114<br>2.6% | 34/114<br>29.8% | 34/114<br>29.8% | 40/114<br>35.1% |

Abbreviations: S=streptomycin, E=ethambutol, Z=pyrazinamide, Km=kanamycin, Am=amikacin, Cm=Capreomycin, Fq=fluoroquinolone, Thio=thioamide, PAS=para-aminosalicylic acid

**Appendix Table S5:** Likelihood scores for different substitution models calculated with Jmodeltest 2.1 and statistical model selection based on Akaike and Bayesian Information Criteration (AIC and BIC). Best model is assumed to have the lowest criteration value. Shown are the top 10 AIC models. Substitution model used for Bayesian inference marked in bold.

| Subst. model | -lnL | AIC | Δ AIC<br>(AIC ranking) | BIC | Δ BIC<br>(BIC ranking) |
| --- | --- | --- | --- | --- | --- |
| **GTR** | 8837.6437 | **18567.2875** | **0.0 (1)** | **21041.0025** | **7.0748 (2)** |
| GTR+I | 8837.6747 | 18569.3494 | 2.0619 (2) | 21048.6109 | 14.6832 (5) |
| GTR+G | 8838.9842 | 18571.9684 | 4.6809 (3) | 21051.2299 | 17.3022 (6) |
| GTR+I+G | 8839.0077 | 18574.0153 | 6.7278 (4) | 21058.8233 | 24.8955 (8) |
| TPM1uf | 8845.426 | 18576.852 | 9.5645 (5) | 21033.9277 | 0.0 (1) |
| TPM1uf+I | 8845.4446 | 18578.8891 | 11.6016 (6) | 21041.5113 | 7.5836 (3) |
| TPM1uf+G | 8846.7354 | 18581.4709 | 14.1834 (7) | 21044.093 | 10.1653 (4) |
| TPM1uf+I+G | 8846.7697 | 18583.5395 | 16.252 (8) | 21051.7081 | 17.7804 (7) |
| **SYM** | 8860.6478 | 18607.2955 | 40.008 (9) | 21064.3712 | 30.4435 (9) |
| SYM+I | 8860.6826 | 18609.3652 | 42.0777 (10) | 21071.9874 | 38.0596 (12) |

## Additional data

34 Resistance targets (in 28 genes) and molecular markers considered as resistance predictor. Phylogenetic variants in 34 resistance associated target genes.

DST phenotypes and polymorphisms in resistance and compensatory genes found in 705 MDR-MTBC strains from Karakalpakstan, Uzbekistan and Samara, Russia and 19 CAO-Beijing strains from Germany. Genotypic classification, transmission indexes, Accession numbers.38 Central Asian outbreak (CAO) specific SNPs with annotations.

Author contributions
S.N., M.M., T.W. H.C. and P.S. designed the study. M.M., M.B., H.C., S.F., J.P.R., U.N., R.D., S.B., S.G.,V.N., S.A., and T.W. analyzed data e.g. classical mycobacteriology, performed population genetic, phylogenetic and statistical analysis. T.A.K. performed whole genome sequencing and variant calling. All authors analyzed the data and contributed to data interpretation and manuscript writing

